# Mechanical activation of VE-cadherin stimulates AMPK to increase endothelial cell metabolism and vasodilation

**DOI:** 10.1101/2024.05.09.593171

**Authors:** Nicholas M. Cronin, Logan W. Dawson, Kris A. DeMali

## Abstract

Endothelia cells respond to mechanical force by stimulating cellular signaling, but how these pathways are linked to elevations in cell metabolism and whether metabolism supports the mechanical response remains poorly understood. Here, we show that application of force to VE-cadherin stimulates liver kinase B1 (LKB1) to activate AMP-activated protein kinase (AMPK), a master regulator of energy homeostasis. VE-cadherin stimulated AMPK increases eNOS activity and localization to the plasma membrane as well as reinforcement of the actin cytoskeleton and cadherin adhesion complex, and glucose uptake. We present evidence for the increase in metabolism being necessary to fortify the adhesion complex, actin cytoskeleton, and cellular alignment. Together these data extend the paradigm for how mechanotransduction and metabolism are linked to include a connection to vasodilation, thereby providing new insight into how diseases involving contractile, metabolic, and vasodilatory disturbances arise.

## INTRODUCTION

All cells in the human body experience forces. These forces are sensed by cell surface receptors and are propagated to the cell interior. To counter these applied forces, cells reinforce their adhesion to neighboring cells and to their actin cytoskeletons (Liu *et al*., 2010; Chanet and Martin, 2014). Reinforcement requires increases in protein synthesis, transport, actomyosin contractility, and actin polymerization— all of which are processes that require energy (Daniel *et al*., 1986; Bernstein and Bamburg, 2003).

How cells derive the energy they need to support reinforcement of the cell-cell adhesions and the actin cytoskeleton is an area of active investigation. In response to increases in tension, modifications to the enzymes and allosteric modulators involved in energy metabolism are altered (Dawson, Cronin and DeMali, 2023). In epithelial cells, E-cadherin senses external forces, the tension is propagated through the protein to produce conformational changes on the cytoplasmic domain (Borghi *et al*., 2012). These conformational changes are believed to allow for recruitment of binding partners and activation of signal transduction cascades. One such signal transduction cascade is initiated by AMPK-activated protein kinase (AMPK), a master regulator of cell metabolism. AMPK is activated and recruited to E-cadherin in response to application of external forces (Bays *et al*., 2014). Active AMPK has two effects: it stimulates a signal transduction cascade that culminates in robust changes in the actin cytoskeleton allowing the cell to resist increased external tension. Additionally, it calls for an increase in glucose uptake through glucose transporter-1 which is required for the generation of ATP (Salvi *et al*., 2021). The increased ATP fuels the alterations in protein synthesis, transport, actomyosin contractility, and actin polymerization needed for the cell to counter the applied tension.

Much of the work accomplished to date has been performed in epithelial cells. The broader applicability of this paradigm has yet to be rigorously tested. Endothelial cells offer a promising model for testing this possibility, as they share a common feature with epithelial cells: both express a force-sensing cell surface cadherin, namely vascular endothelial cadherin (VE-cadherin) in endothelial cells(Barry et al., 2015)(Conway and Schwartz, 2015). Additionally, AMPK is activated in endothelial cells in response to force (Li, Sun and Carmeliet, 2019). However, there are additional levels of complexity in endothelial cells which cast doubt as to whether mechanics and metabolism are coupled in the same manner observed in epithelial cells. Key among these is that force sensing is more intricate in endothelial cells. VE-cadherin is part of a mechanosensory complex, including platelet endothelial cell adhesion molecule 1 (PECAM-1) and vascular endothelial growth factor receptor 2 (VEGFR2). Additionally, VE-cadherin only bears force when present in adherens junction (Conway *et al*., 2013) This contrasts with E-cadherin, which bears force when it is on the cell surface and irrespective of its presence in adherens junctions (Borghi *et al*., 2012). Finally, while both endothelial and epithelial cells heavily rely upon glycolysis to meet their energetic demands, endothelial cells have fewer mitochondria and do not employ oxidative phosphorylation to the same extent as epithelial cells to generate energy (Dawson, Cronin and DeMali, 2023). These observations raise the question as to how AMPK is activated in response to external forces in endothelial and the consequences of its activation.

Here we present evidence VE-cadherin transduces mechanical cues to the cell interior where liver kinase B1 (LKB1) stimulates AMPK. Active AMPK phosphorylates endothelial nitric oxide synthase (eNOS), thereby coupling VE-cadherin stimulated mechanotransduction and vasodilation. VE-cadherin stimulated AMPK calls for increased glucose uptake and metabolism, to provide the energy necessary for reinforcement of the actin cytoskeleton and cadherin adhesion complex and endothelial cell alignment. Taken together this work extends the paradigm for how mechanotransduction and metabolism are linked to include a connection to vasodilation.

## RESULTS

### VE-Cadherin is required for shear stress induced activation of AMPK

Cells respond to mechanical cues from their environment by reinforcing their cytoskeleton and adhesions, a process that consumes considerable energy (Daniel *et al*., 1986; Bernstein and Bamburg, 2003; Choi and Helmke, 2008). The mechanisms by which this energy is produced remain poorly understood. However, in epithelial and endothelial cells, AMP-activated protein kinase (AMPK), a master regulator of cell metabolism, is activated in response to fluid shear stress (Young *et al*., 2009; Bays *et al*., 2017), suggesting its involvement. To explore the role of AMPK, we monitored its activation in human umbilical vein endothelial cells (HUVECs) and bovine aortic endothelial cells (BAECs) subjected to varying durations and intensities of shear stress. Activation of AMPK was assessed by examining phosphorylation of AMPK in its activation loop using a Thr172 phospho-specific antibody, an approach that we and other groups have shown mirrors AMPK kinase activity and phosphorylation of its substrates (Bays et al,2019). Consistent with previous reports, AMPK was activated in response to shear stress (Young et al, 2009). Maximal activation occurred with application of 12 dynes/cm^2^ of shear stress and after 5 minutes (Fig S1A-B) in HUVECs and (Fig S1D-E) in BAECs.

The activation of AMPK in response to shear stress suggested that VE-cadherin, a major force transducing protein on the surface of endothelial cells, might be necessary for this process. To examine the role of VE-cadherin in shear stress stimulated AMPK activation, cells were incubated with BV9, a VE-cadherin function blocking antibody, or DMSO as a control. After treatment, cells were either left undisturbed (-) or subjected to 12 dynes/cm² of shear stress (+) for durations ranging from 5 to 20 minutes. AMPK was maximally activated with a 3.5 ±1.1-fold increase after 5 minutes of applied shear stress. AMPK activation was not increased in cells pre-incubated with the VE-cadherin function blocking antibody (Fig. 1A-B).

**Figure 1:**
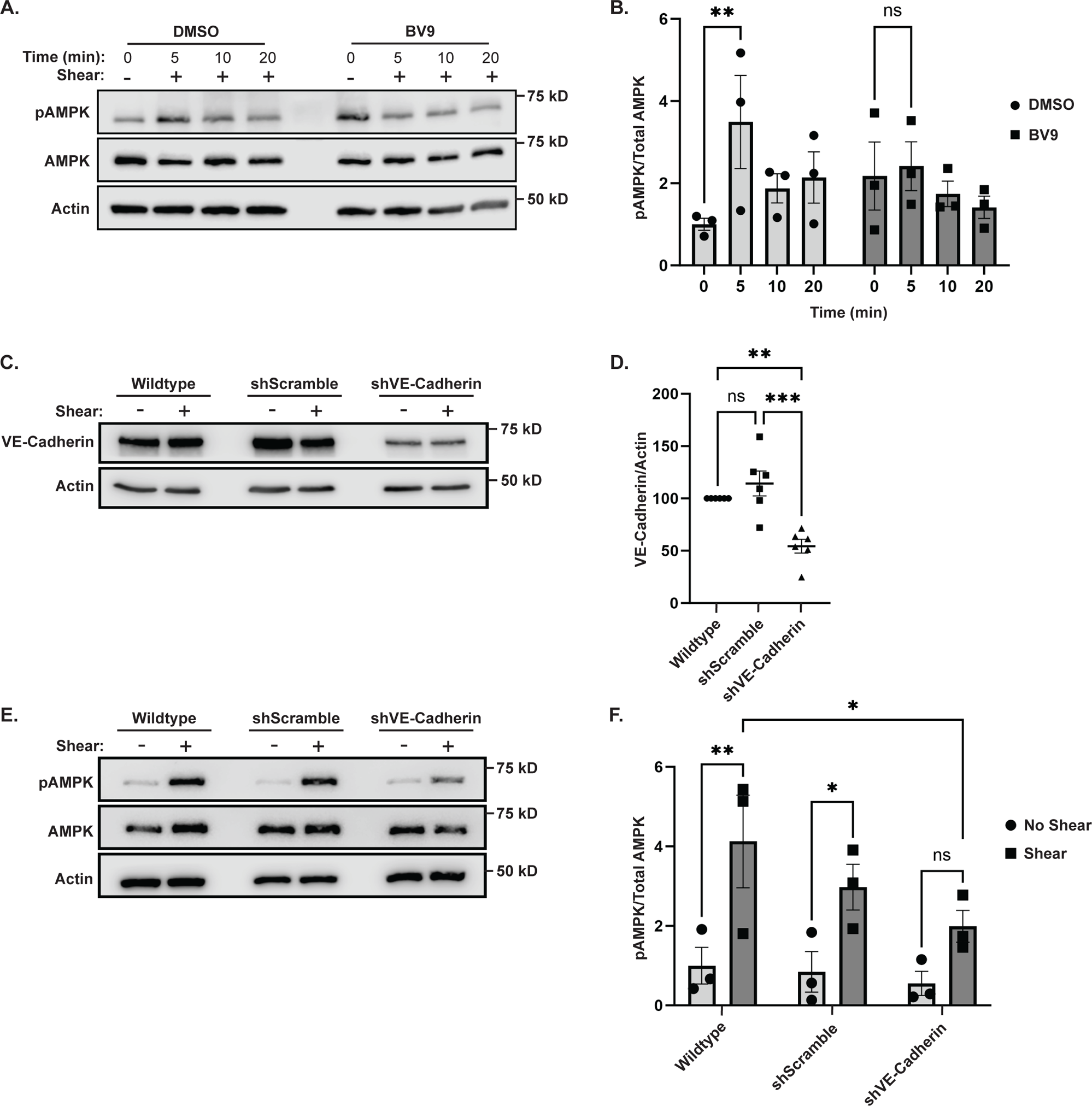
VE-Cadherin is required for shear-induced AMPK activation. (A-B). Shear stimulated AMPK activation in the presence of VE-cadherin function blocking antibodies. HUVECs were left resting (-) or exposed to shear stress (+) for the indicated times in the presence or absence of BV9, a VE-cadherin function blocking antibody, or DMSO as a control. AMPK activation was examined by immunoblotting total cell lysates with an antibody that recognizes AMPK phosphorylation in its activation loop (pAMPK) or actin as a loading control. The blots were stripped and probed with antibodies that report on total AMPK levels (AMPK). The graph in B depicts the ratios of phosphorylated AMPK to total AMPK; the data are mean ±s.e.m., n =3 biologically independent samples. **p<0.01, (two-way ANOVA). (C and D) VE-cadherin inhibition. VE-cadherin expression was inhibited by treating HUVECs with lentiviruses encoding shRNAs targeting human VE-cadherin (shVE-Cadherin),or a scrambled sequence (shScramble). Wildtype parental cells were employed as a control. Total cell lysates were immunoblotted with antibodies against VE-cadherin to show the level of inhibition or actin as a loading control Representative western blot images are shown (C). The graphs in (D) depict the level of VE-cadherin normalized to total protein levels. The data are mean ±s.e.m n=6 biologically independent samples. **P<0.01 (One-way ANOVA, with Dunnett’s test). ns=not significant. (E and F) Shear stress induced AMPK activation in cells with depressed VE-cadherin levels. AMPK activation was examined as described above in A and B in wildtype, shScramble, or shVE-cadherin cells. The data are mean ±s.e.m n=6 biologically independent samples. *P<0.05, **P<0.01 (Two-way ANOVA, with Tukey’s comparison), ns=not significant.

As a second measure, we suppressed VE-cadherin expression by using RNA interference. To this end, lentiviruses encoding shRNAs against VE-Cadherin (shVE-cadherin) or a scramble sequence (shScramble) as a control were generated and transduced into HUVECs. Seventy-two hours after viral infection, VE-Cadherin expression was decreased by 54±11% (Fig. 1C-D) when compared to the parental HUVECs (Wildtype). To determine if AMPK was activated when VE-cadherin levels are depressed, the cells were incubated with and without 12 dyne/cm^2^ of shear stress. AMPK phosphorylation was significantly reduced in the cells depleted of VE-Cadherin (Fig. 1E-F), confirming the essential role of VE-Cadherin in facilitating AMPK activation in response to shear stress.

### LKB1 is required for mechanical activation of AMPK and eNOS

AMPK can be targeted by three known kinases: LKB1, CaMKKβ, and TAK1. Previous work implicated CaMKKβ in AMPK activation (Hurley *et al*., 2005). However, inhibiting CaMKKβ with the CaMKKβ inhibitor STO-609 did not prevent shear stress induced AMPK activation (Fig. S2A-B). This led us to explore whether LKB1 might be responsible for AMPK activation under these conditions. To test the possibility that LKB1 facilitates AMPK activation in response to shear stress, we generated lentiviruses encoding shRNAs against LKB1 and infected HUVECs with the viral particles. Lentiviral application produced a 65±15% decrease in LKB1 levels (Fig 2A-B). Furthermore, this level of inhibition significantly decreased shear stress induced AMPK activation (Fig. 2C-D).

**Figure 2:**
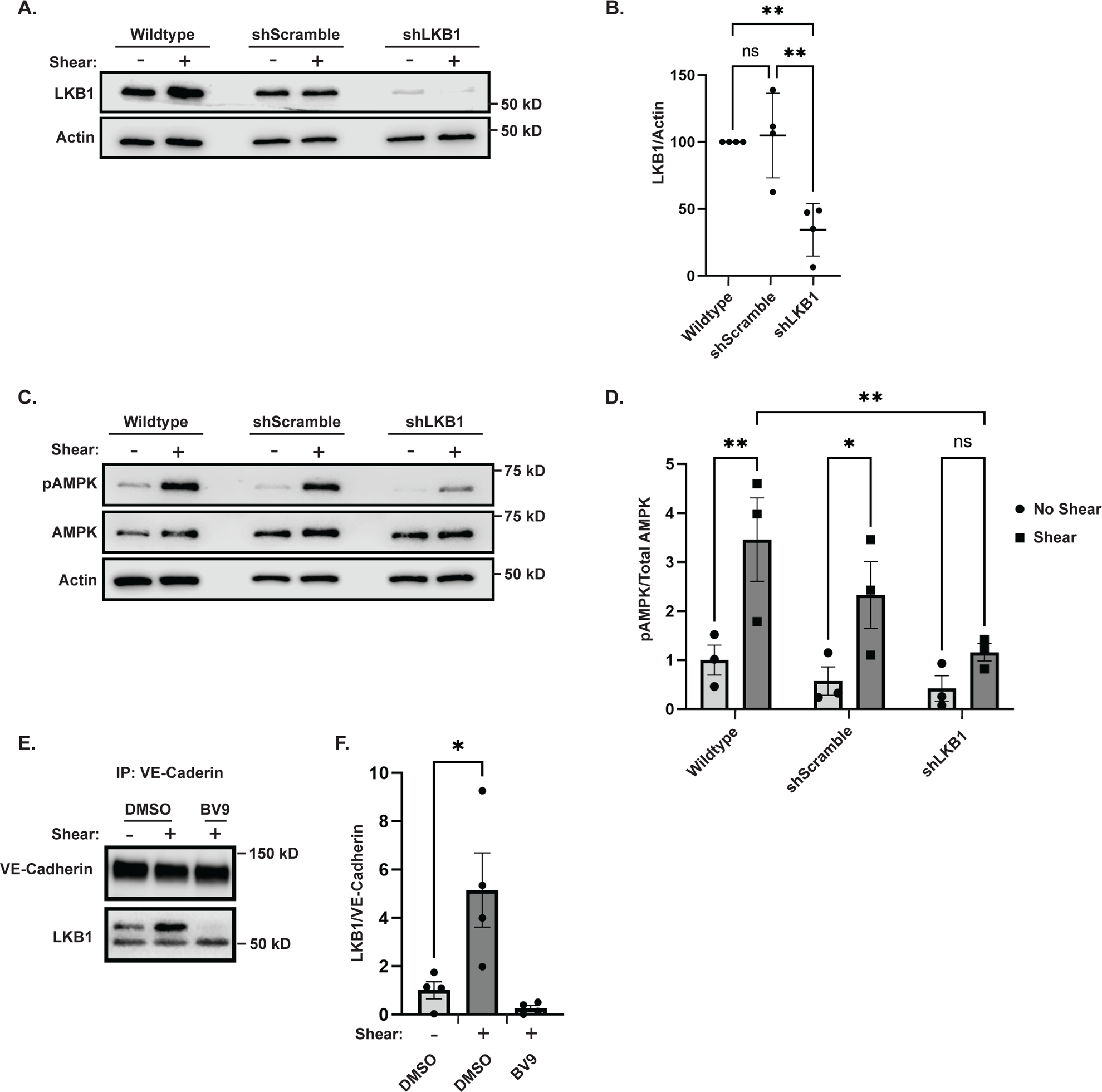
LKB1 is the upstream kinase responsible for shear-induced AMPK activation. (A) LKB1 inhibition using RNA interference. LKB1 expression was inhibited by treating HUVECs with lentiviruses encoding shRNAs targeting human LKB1 (shLKB1), or a scramble sequence (shScramble). Wildtype parental cells were employed as a control. Total cell lysates were immunoblotted with antibodies against LKB1 to show the level of inhibition or actin as a loading control. Representative western blots are shown. (B) The graph depicts the LKB1 levels normalized to the total protein levels; the data are mean ±s.e.m., n =4 biologically independent samples. **p<0.01 (One-way ANOVA, with Dunnett’s test). (C) Shear stress stimulated AMPK activation in cells depleted of LKB1. AMPK activity was monitored as described in Figure 1A. Representative immunoblots are shown. The graph in (D) depicts the ratio of phosphorylated AMPK to total protein. The data are mean ±s.e.m., n=3 biologically independent samples. ns=not significant, *P<0.05, **P<0.01 (RM two-way ANOVA, with Tukey’s multiple comparison test). (E-F) Shear induced LKB1 coimmunoprecipitation with VE-Cadherin. Cells were preincubated with DMSO or BV9 for 1 hour and then were left resting (-) or exposed to shear stress (+) for 5 minutes. VE-Cadherin was immunoprecipitated, and the co-precipitation of LKB1 was examined. The ratio of LKB1 to VE-Cadherin can be seen in the graph (F). Data are mean ±s.e.m., n=4 biologically independent samples, *P<0.05, (One-way ANOVA, with Dunnett’s test).

Our observation that LKB1 is required for shear stress induced AMPK and the observation that LKB1 is localized to the plasma membrane in response to shear stress (Bays *et al*., 2017; Dogliotti *et al*., 2017), suggested the possibility that LKB1 might be recruited to the cadherin adhesion complex. To test this possibility, LKB1 was recovered from cells that had been left resting or subjected to shear stress. VE-cadherin co-immunoprecipitation was examined by immunoblotting. Exposure to shear showed increased VE-cadherin binding to LKB1. Furthermore, the co-immunoprecipitation was lost upon pre-incubation of the cells with Bv9 function blocking VE-cadherin antibody or inhibition of VE-cadherin expression (Fig. 2E-F). These data suggest that LKB1 recruitment to VE-cadherin upon exposure to shear stress is required for AMPK activation.

AMPK phosphorylates many downstream targets (Mihaylova and Shaw, 2011). One target of AMPK is endothelial nitric oxide synthase (eNOS) (Morrow *et al*., 2003; Thors, Halldórsson and Thorgeirsson, 2011). When activated, eNOS generates the potent vasodilator nitric oxide a molecule critical for proper endothelial function (Durand and Gutterman, 2013). Thus, we next determined if VE-cadherin and AMPK was required for eNOS activation. To address this possibility, cells were preincubated with BV9 (a VE-cadherin function blocking antibody) or DMSO and left at rest or subjected to 12 dynes/cm^2^ shear stress. eNOS phosphorylation was assessed by blotting lysates with a phospho-specific antibody that targets Ser1177, a known AMPK phosphorylation site (Thors, Halldórsson and Thorgeirsson, 2011). After 10 minutes of shear stress, phosphorylation of Ser1177 was increased by 2.0±0.5-fold in the DMSO treated cells (Fig. 3A-B). In contrast, preincubation of cells with BV9 prevented a significant increase in eNOS phosphorylation (Fig 3A-B). Since LKB1 is required for VE-cadherin stimulation of AMPK, we explored if LKB1 was required for eNOS activation triggered by shear stress. Transient inhibition of LKB1 using shRNAs decreased eNOS stimulation in response to shear stress (Fig. 3C-D). To further assess the role of VE-cadherin and LKB1 in the mechanical activation of eNOS, we examined eNOS localization. eNOS myristylation and recruitment to the plasma membrane regulates its activity (Sakoda *et al*., 1995; Shaul *et al*., 1996), but the mechanism of this recruitment is not well defined. Since VE-Cadherin is highly expressed in the plasma membrane of endothelial cells, and stimulates AMPK to phosphorylate eNOS, we explored how VE-cadherin influences eNOS localization to adherens junctions. HUVECs were left resting or subjected to 12 dyne/cm^2^ of shear stress and eNOS enrichment at cell-cell junctions was examined using confocal microscopy. eNOS was enriched in VE-cadherin cell-cell junctions in response to shear stress with a Pearson’s Correlation Coefficient of 0.54 ± 0.1 (Fig 3E-G). This localization was dependent upon VE-cadherin, LKB1, and AMPKα1 as their depletion prevented eNOS localizations (Fig 3E-G). Taken together, these findings indicate eNOS localization to cell-cell junctions requires not only VE-cadherin, but also LKB1-mediated activation of AMPK. These data reveal that mechanical cues transduced by VE-cadherin are required eNOS activation and function.

**Figure 3:**
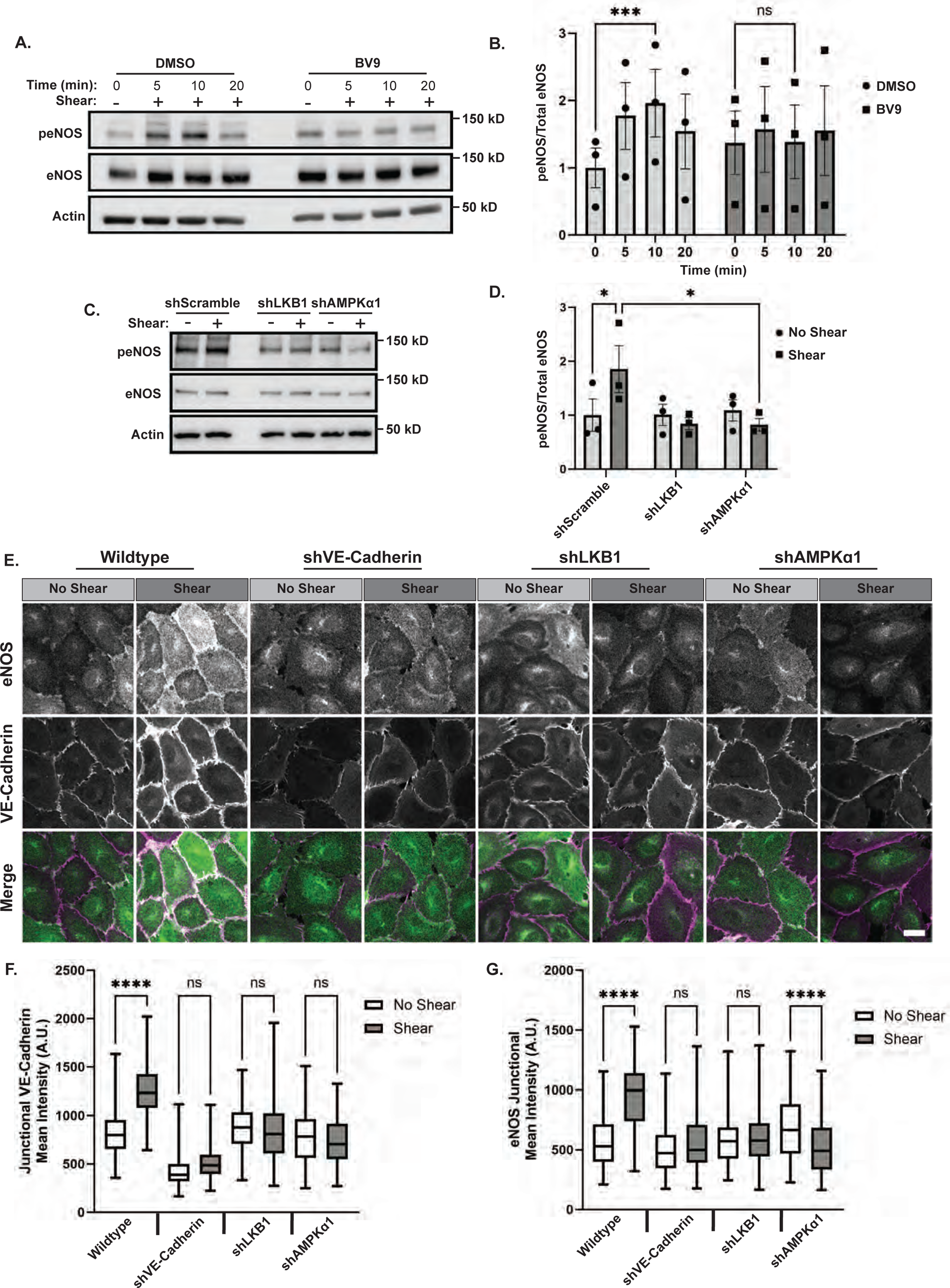
VE-Cadherin stimulated AMPK is required for eNOS membrane localization and activation. (A) eNOS activation in the presence of VE-cadherin inhibitors. HUVECs were left untreated (DMSO) or treated with 5μg BV9 and left resting conditions (-) or exposed to12 dyne/cm2 orbital shear (+) for indicated times in minutes. (B) The graphs to the right depict the ratio of phosphorylated protein to total protein; the data are mean ±s.e.m., n =3 biologically independent samples. **p<0.01, (RM two-way ANOVA, with Tukey’s test). (C-D) eNOS activation when VE-Cadherin stimulated LKB1 is disrupted. Wildtype HUVECs or those expressing shRNA against LKB1 (shLKB1) or AMPKα1 (shAMPKα1) were left under restring (no shear) or exposed to shear. The cells were lysed and probed with an eNOS Ser 1177 phospho-specific antibody. The ratio of phosphorylated to total protein, can be seen in the quantification in graph (D). The data are mean ±s.e.m., n =3 biologically independent samples. **p<0.01, (RM two-way ANOVA, with Tukey’s multiple comparison test). (E-G) eNOS localization when VE-cadherin stimulated AMPK is disrupted. Wildtype HUVECs or those expressing shRNAs against VE-Cadherin (shVE-Cadherin), LKB1 (shLKB1) or AMPKα1 (shAMPKα1) were left resting (no shear) or exposed to shear for 30 minutes. The cells were fixed and stained with antibodies against eNOS (green) or VE-Cadherin (magenta) and examined by confocal microscopy. Representative images are shown in E, and an average Pearson correlation coefficient ± STD is reported beneath the images. Scale bars = 20 μm. The graphs beneath the images represent he average corrected fluorescence intensity of VE-Cadherin (F) or eNOS (G) in 100 junctions per condition and cell type. The data are represented as box and whisker plots with median 10th, 25th, 75th and 90th percentiles and min and max shown, n = 100 over 2-3 fields of view. ****p<0.001 (One-way ANOVA, with Tukey’s multiple comparison test).

### VE-Cadherin-mediated AMPK activation is required for actin cytoskeletal reinforcement

Shear stress stimulates reinforcement of the actin cytoskeleton and cell-cell adhesions. The requirement for AMPK for these events is unknown. To test a role for AMPK in junctional actin reinforcement, shear stress (12 dynes/cm^2^) or no shear stress was applied to confluent cultures of parental HUVECs (wildtype) or HUVECs expressing shRNAs against LKB1 (shLKB1), AMPK (shAMPKα1) or a scramble sequence (shScramble) as a control. Junctional morphology and enrichment of the actin cytoskeleton were monitored via immunofluorescence. Under no shear stress conditions, both wildtype and shScramble cells displayed a fenestrated pattern of VE-Cadherin staining, which shifted to a linear orientation upon application of shear stress. (Fig 4A). Additionally, VE-cadherin was enriched in junctions in response to shear stress. This is consistent with previous reports on reinforcement of cell-cell junctions (Fig 4A-B). The actin cytoskeleton was also reinforced when shear was applied (Fig. 4A, C). However, these structural reinforcements were lost when either LKB1 or AMPKα1 was depleted (Fig. 4A-C). Thus, VE-cadherin mediated changes in LKB1 and AMPKα1 are required for reinforcement of the actin cytoskeletal and adherens junctions.

**Figure 4:**
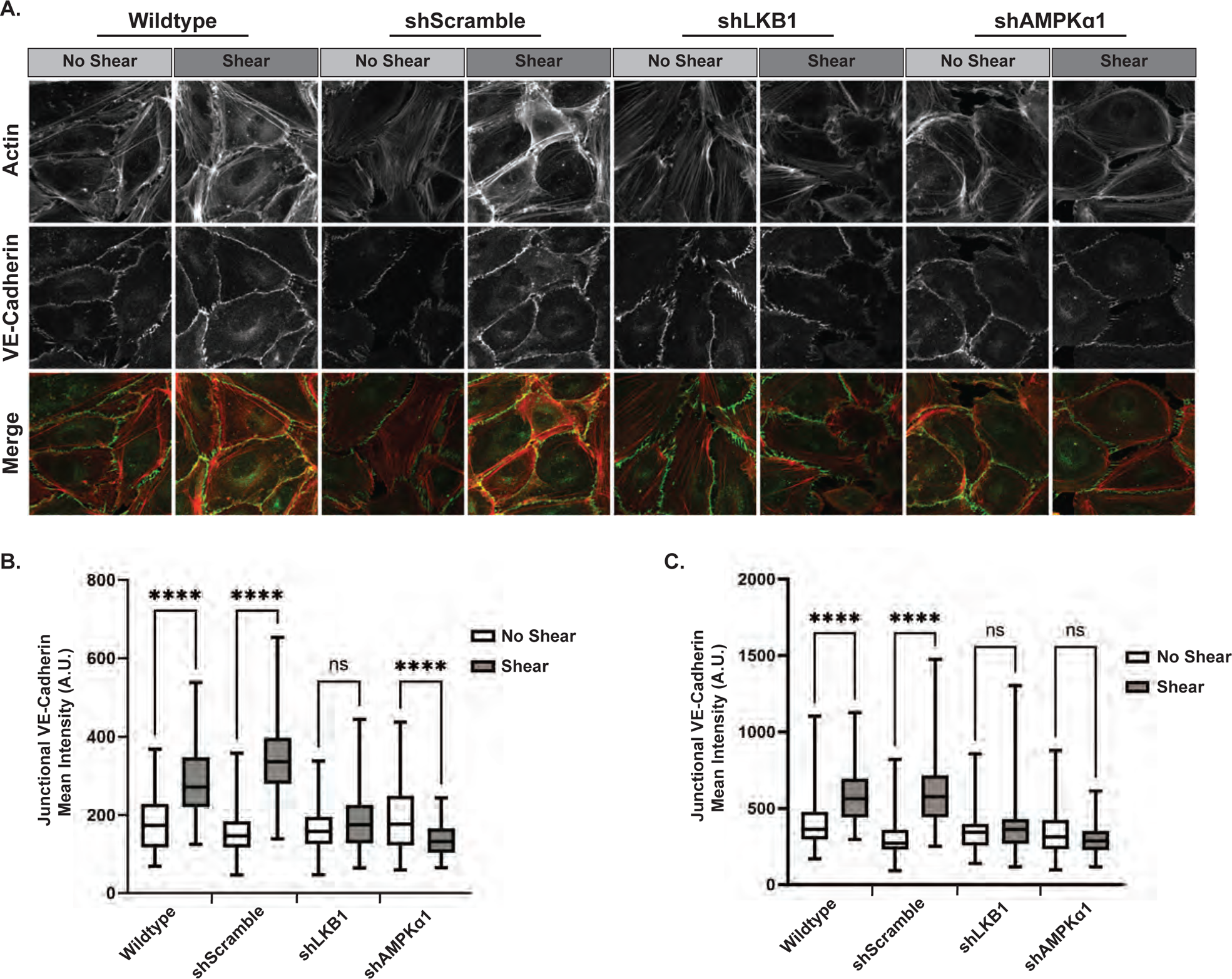
LKB1 and AMPKa1 are required for shear-induced junctional reinforcement. (A-C) Effect of AMPK inhibition of actin cytoskeletal reinforcement. Wildtype HUVECs or those expressing shRNAs against LKB1 (shLKB1), AMPKα1 (shAMPKα1), or a scrambled sequence (shScramble) were left resting (no shear) or exposed to shear for 30 minutes. The cells were fixed and stained an antibody against VE-Cadherin (green) or Phalloidin conjugated to Alexa594 to visualize actin (red). Cells were examined by confocal microscopy. Representative images are shown in A. Scale bars = 20 μ m. The graphs beneath the images represent the average corrected fluorescence intensity of actin (B) or VE-Cadherin (C) in 100 junctions per cell type. The data are represented as box and whisker plots with median 10th, 25th, 75th and 90th percentiles and min and max shown, n = 100 over 2-3 fields of view. ****p<0.0001 (One-way ANOVA, with Tukey’s multiple comparison test).

### VE-Cadherin-mediated AMPK calls for increased metabolism

Our observations that VE-cadherin increases AMPK activity in response to mechanical cues prompts investigation into the consequences of this activation. The cellular response to mechanical cues requires elevations in enzymatic activity, actin polymerization, and actomyosin contractility, all of which are energy-intensive processes necessary for cytoskeletal rearrangement and adhesion growth (Dawson, Cronin and DeMali, 2023). Given that glucose is the preferred energy source for endothelial cells, we hypothesized that the increase in AMPK activity due to VE-cadherin could enhance glucose metabolism. To begin to test this, glucose uptake was examined in HUVECs subjected to shear stress. For this, shear stress was applied to HUVECs, and the uptake of a fluorescently labelled, non-hydrolyzable 2-deoxyglucose (2-NBDG) analog was monitored. This analysis revealed that glucose uptake was increased 3.11-fold in response to application of shear stress (Fig 5A-B). This uptake specifically involved the most predominant glucose transporter in epithelia, glucose transporter 1 (GLUT1), as pre-treatment with WZB117, an inhibitor of GLUT1, blocked this effect (Fig. 5B). In contrast, preincubation of cells with 3PO, a phosphofructokinase-2 inhibitor that interferes with glucose metabolism had no effect on shear-induced glucose uptake. To determine if this elevation in glucose uptake required VE-cadherin force transduction, cells were preincubated with a VE-cadherin function blocking antibody or Blebbistatin, a myosin II inhibitor. Both prevented glucose uptake, further suggesting that force transmission is required for elevated. These analyses revealed that VE-cadherin is required for the elevated glucose uptake that was observed in response to application of shear stress (Fig. 5B).

**Figure 5:**
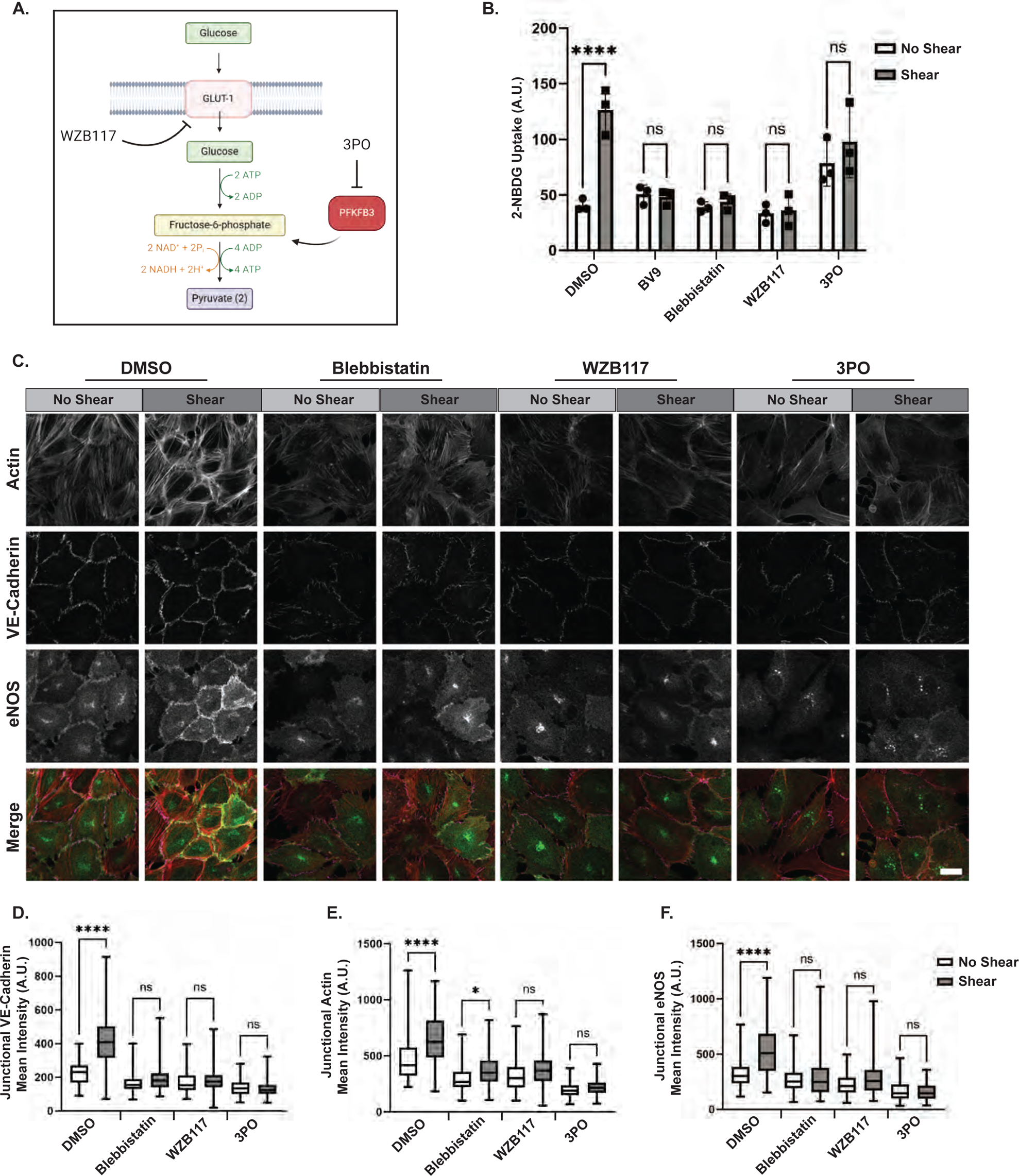
**(A) Illustration of glycolytic inhibitors and their targets**. (B) Mechanical or metabolic disruption and its effects on shear induced glucose uptake. HUVECs were switched to glucose free media and were pretreated with DMSO or inhibitors against VE-Cadherin (BV9), myosin II (Blebbistatin), GLUT1 (WZB117), or phosphofructokinase fructose-bisphosphate 3 (3PO). 2-NBDG was added to cells under no shear or shear conditions for 1 hour. Fluorometric analysis of the cell lysate was performed and the value of 2-NBDG uptake is quantified as the mean ±s.e.m., n=3 biologically independent samples. ns=not significant, ****P<0.001 (RM two-way ANOVA, with Tukey’s multiple comparison test). (C) The effect of metabolic inhibitors on reinforcement of the actin cytoskeleton and cadherin adhesions. HUVECs pretreated with DMSO or inhibitors against myosin II (Blebbistatin), GLUT1 (WZB117), or phosphofructokinase fructose-bisphosphate 3 (3PO). Cells were then left under static (no shear) or exposed to shear stress (shear) conditions for 30 minutes. Cells were fixed and stained with Alexa 594-Phalloidin (red) and antibodies against VE-Cadherin (magenta) and eNOS (green). Representative images are shown in A. Scale bar, 20 μm. The graphs represent the average corrected fluorescence intensity of VE-Cadherin (D), eNOS (E), or actin (F) in 100 junctions per condition. The data are represented as box and whisker plots with median 10th, 25th, 75th and 90th percentiles and min and max shown, n = 100 over 2-3 fields of view. ****p<0.001 (One-way ANOVA, with Dunnett’s multiple comparison test.

### VE-Cadherin-mediated AMPK fuels cellular reinforcement and alignment

Having established VE-cadherin force transmission stimulates glucose uptake, we next explored why glucose is crucial for cellular processes. Previous work indicates that elevations in glucose metabolism are required for a whole array of cellular responses (Pi, Xie and Patterson, 2018). To test the possibility that glucose was required for increased reinforcement of cellular adhesions and the cytoskeleton, cells were treated with either WBZ117, a small molecule inhibitor of the glucose transporter GLUT1, or 3PO, a small molecule inhibitor of PFKFB3, which inhibits glycolytic flux. The cells were then left under no shear or shear conditions and examined by confocal microscopy. A loss of junctional linearization, enrichment of VE-cadherin, and actin cytoskeletal reinforcement was observed when GLUT1 or glycolysis was inhibited (Fig. 5C-E). Interestingly, another consequence preventing glucose uptake or glycolysis was a loss of eNOS localization to cell-cell junctions (Fig. 5C,F). Thus, elevations in glucose uptake and metabolism are required for fortification of the cytoskeleton and adhesion of cells.

A common *in vitro* and *in vivo* physiological response of endothelial cells is to align parallel to the direction of shear stress, an event that is critical in the protection against the development of atherosclerosis and inflammation (Immanuel and Yun, 2023). To test if the VE-cadherin activated metabolism we studied herein is important for endothelial cell alignment, HUVECs were left resting or subjected to shear stress for 48 hours. The cells were stained to visualize actin stress fiber alignment and nuclei orientation. Alignment was assessed by measuring the Feret angle of the major axis (L_major_) of the cells with respect to the direction of the flow. We found that 77.5±5.1% of DMSO treated cells exposed to shear were aligned. However, disruption of mechanotransduction either by non-muscle myosin II inhibition with blebbistatin or VE-Cadherin disruption using BV9 significantly reduced cellular alignment (Fig. 6A-B). Inhibition of glucose uptake or glycolysis also produced decreased alignment (Fig. 6A-B). We sought to determine if the loss of alignment was a consequence of decreased elongation by measuring the ratio between the major (L_major_) and minor (L_minor_) axis of the cells. All the inhibitors prevented cellular elongation (Fig. 6C). Collectively, these findings suggest that shear stress induces VE-cadherin to activate LKB1, which then activates AMPK. This activation of AMPK leads to increased glucose uptake, fueling the structural reinforcement necessary for cells to align in response to mechanical forces.

**Figure 6:**
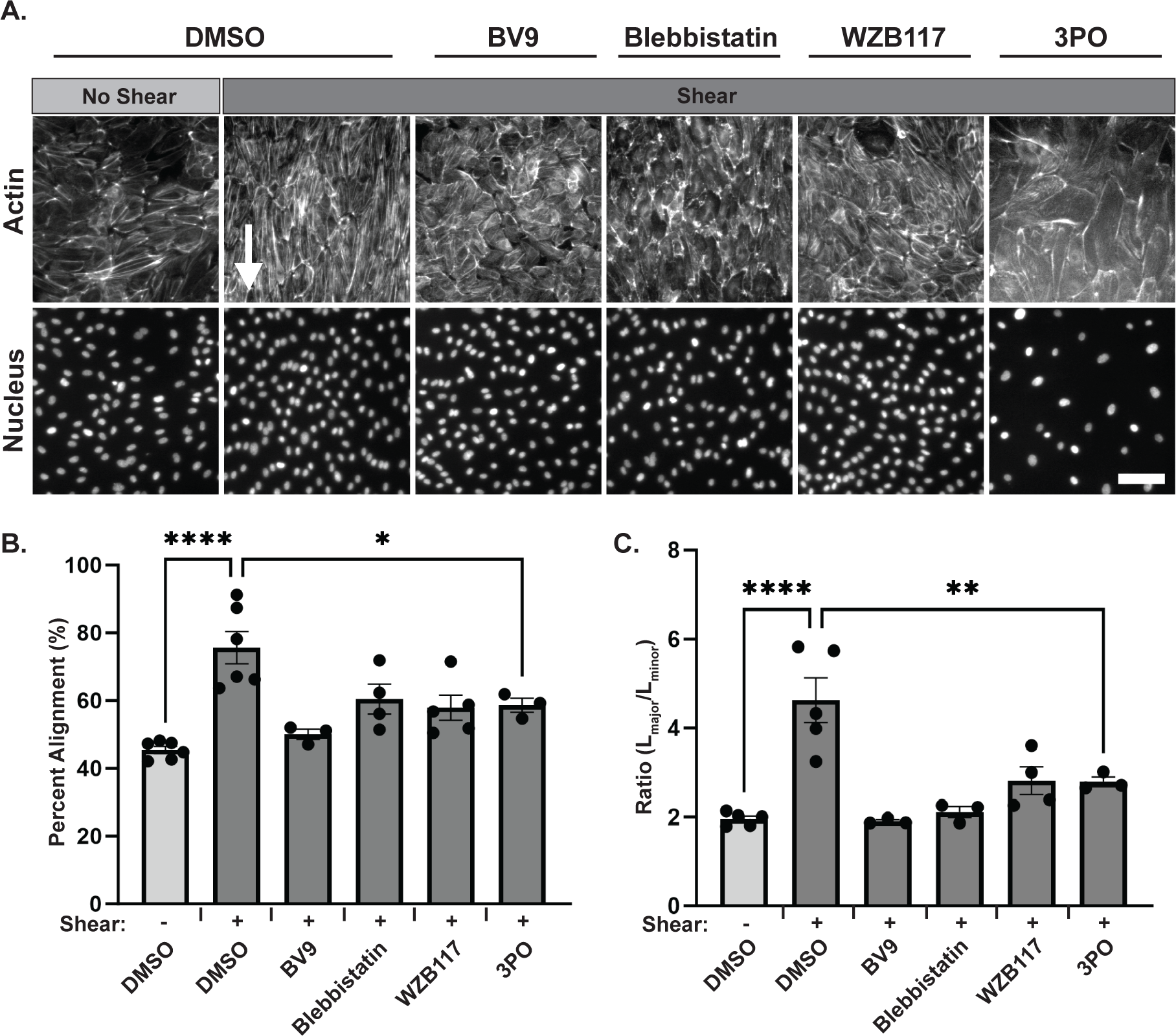
Increased metabolism supports endothelial alignment. (A) The effects of metabolic inhibitors on endothelial alignment. HUVECs pretreated with DMSO or inhibitors against VE-Cadherin homophilic binding (BV9), myosin II (Blebbistatin), GLUT1 (WZB117), or phosphofructokinase fructose-bisphosphate 3 (3PO). Cells were then left under static conditions (no shear) or exposed to shear stress (shear) conditions for 48 hours. Media and inhibitors were reapplied after 24 hours. Cells were fixed and stained with Alexa 594-Phalloidin and DAPI to visualize actin and the nucleus respectively. Representative images are shown in A. Scale bar, 40 μm. Alignment was assessed based on whether the major axis of the cell and its Feret angle was within ±45° of the direction of flow and is quantified in B. Elongation was assessed by comparing the ratio of the length of the major axis (L¬major) with the width of the minor axis (Lminor) and is quantified in C. >250 cells across 3 fields of view per condition were assessed for alignment and 100 cells across 3 fields of view per condition were assessed for elongation. The average percent alignment and elongation is plotted as the mean ±s.e.m., n=3-5 biologically independent experiments. *P<0.05, **P<0.01, ***P<0.001, ****P<0.0001 (One-way ANOVA, with Dunnett’s multiple comparison test).

## DISCUSSION

The response of cells to mechanical cues requires increases in protein synthesis, transport, actomyosin contractility, and actin polymerization needed for the cell to counter the applied tension (Dawson, Cronin and DeMali, 2023). All these processes require energy. In epithelia, increased tension is propagated across E-cadherin and signals for increased AMPK activity which stimulates a downstream signal transduction cascade that culminates in increased contractility (Bays *et al*., 2017). Here we report the existence of a similar pathway in endothelia. Indeed, shear stress stimulated activation of AMPK. This activation was dependent on VE-cadherin and produced elevations in eNOS activity. These findings not only confirm the linkage between cell mechanics and metabolism, as observed in epithelial cells, but also extend this relationship to include eNOS activity and vasodilation, further expanding the emerging paradigm of how mechanical and metabolic processes are interconnected.

What is the increase in metabolism needed for? Numerous independent studies demonstrated that actin cytoskeleton is a major energetic demand in cells (Daniel *et al*., 1986; Bernstein and Bamburg, 2003; Gómez-Escudero *et al*., 2019) and that altered cytoskeletal dynamics affect cell metabolism (Dawson, Cronin and DeMali, 2023). We previously reported that increased metabolism in epithelial cells supports cytoskeletal rearrangements and growth of the adhesion complex in response to tension (Salvi *et al*., 2021). Here we find that perturbing force-stimulated metabolism blocks actin cytoskeletal reinforcement, reinforcement of the cadherin adhesion complex, and cellular alignment (Figs 4, 5 and 6). All these events require robust actin cytoskeletal changes and are blocked by inhibitors of cellular metabolism, thereby suggesting a requirement for increased metabolism to support the actin cytoskeleton.

The notion that energy is required for supporting the actin cytoskeleton has recently been challenged (Holland and Gallo, 2023). This study employed a genetically encoded biosensor that indicates its ATP level based on the ratio of ADP to ATP and reported that ATP/ADP ratios remain relatively constant when growth cones are stimulated with nerve growth factor which increases actin dynamics. The authors also present evidence that inhibiting actin cytoskeletal rearrangements has no effect on ATP/ADP ratios. While the biosensors used in this study have provided utility in some settings, they report on ADP/ATP ratios, not ATP concentrations and are susceptible to optical overlap with cellular sources of auto-fluorescence, the latter of which is expected to be high in a growth cone (Lobas *et al*., 2019). Nonetheless, these studies raise the remote possibility that the energy demands for reinforcing the cytoskeleton and adhesion complex, as well as for cellular alignment, might be met by other energy-intensive processes such as protein phosphorylation, transport, and synthesis. Further research is needed to clarify which of these processes are essential when a cell is under tension.

Unlike many other cell types, endothelia can increase vasodilation through the synthesis and release of vasoactive substances, such as nitric oxide, prostacyclin, and endothelium derived hyperpolarizing factor (Durand and Gutterman, 2013). Here we show a 2-fold increase in eNOS phosphorylation was observed in cells under shear stress (Fig 3, S1), corroborating previous findings that shear stress stimulates eNOS activity (Nishida *et al*., 1992; Kolluru *et al*., 2010; Thors, Halldórsson and Thorgeirsson, 2011). Importantly, our results show that this increased eNOS activation requires VE-cadherin (Fig 3), a novel finding not previously reported. To date, shear stimulated increases in eNOS activity are known to occur in response to increased ion channel conductance (Ungvari *et al*., 2001), VEGF treatment (Feliers *et al*., 2005), and thrombin activation (Thors *et al*., 2008) The involvement of VE-cadherin we report here adds a significant new dimension to the understanding of eNOS regulation under shear stress.

In addition to requiring VE-cadherin, shear stress stimulated increases in eNOS phosphorylation required AMPK (Fig 3), and other work demonstrated eNOS is a direct substrate for AMPK (Thors, Halldórsson and Thorgeirsson, 2011). Collectively, these data suggest the existence of a mechanosignaling pathway whereby VE-cadherin stimulated AMPK phosphorylates eNOS to increase NO production and elevate vasodilation. In this scenario, mechanics, metabolism, and vasodilation are linked. The importance of coupling mechanical and metabolic signaling with vasodilation remains speculative. Since vasodilation enhances the flow of blood to areas of the body that lack nutrients, it is tempting to speculate that mechanical signaling is linked to vasodilation to elevate the delivery of these nutrients.

Emerging evidence suggests the links between mechanics, metabolism and vasodilation are much more complex than presented herein. An RNAi screen aimed at identifying cadherin interacting proteins unveiled 400 proteins organized into 17 regulatory hubs with the metabolism hub being the third largest (Toret *et al*., 2014). Additionally, numerous studies demonstrate a complex relationship between Rho kinase, a key mediator of contractility, and the metabolic regulator, AMPK. (Noda *et al*., 2014; Huang *et al*., 2018). Finally, our current studies revealed that inhibition of glycolysis decreases eNOS localization to the plasma membrane, implying that without the generation of glycolytic intermediates, there is less need for increased vasodilation to transport nutrients. This is likely not the only mechanism for coordinating the activities of these interlinked pathways as eNOS interacts and S-nitrosates pyruvate kinase M2, which ultimately decrease substrate flux through the pentose phosphate pathway, thereby decreasing production of NADPH, an important reducing equivalent (Siragusa *et al*., 2019). More work is needed to unravel the complexities of the linkages between mechanics, metabolism, and vasodilation and to better understand their regulation.

## Author contributions

Conceptualization: NC, KAD; Methodology: NC; Validation: NC; Formal analysis: NC, LD; Investigation: NC, LD; Data curation: NC, LD; Writing - original draft: NC, KAD; Writing - review & editing: KAD, LD, NC. Supervision: KAD; Project administration: KAD; Funding acquisition: KAD.

## Methods

### Cell lines

Human Umbilical Vein Endothelial Cells (HUVECs, Lonza Scientific) and Bovine Aortic Endothelial Cells (BAECs, American Tissue Culture Collection) were maintained in MCDB 131 media (SIGMA-Aldrich), supplemented with 5% heat-inactivated FBS, 12ug/ml Bovine Brain Extract (Lonza CC-4098), 10ng/ml hEGF (Sigma E9644), 1ug/ml hydrocortisone (Fisher Scientific), 200ug/ml ascorbic acid (Sigma), and 5% Pen-strep (GIBRO-BRL). All cell lines were used for no more than 6 passages and periodically checked for mycoplasma contamination. The 293FT cells (Invitrogen) are virus-producing cells that are a derivative of 293T cells and were maintained in DMEM supplemented with 10% heat-inactivated FBS, 0.1mM Non-essential Amino Acids (GIBCO-BRL), 6mM L-glutamine, 1mM MEM Sodium Pyruvate, and 1% Pen-Strep, 500 ug/ml Geneticin (GIBCO-BRL).

### Constructs

To inhibit expression of proteins using small hairpin RNAs, the following cDNA sequences were cloned into pLKO.1 which was a gift from David Root (Addgene plasmid # 10878). by Genscript Corporation:

shVE-CADHERIN 5’-CCGGAGATGCAGAGGCTCATGATCTCGAGATCATGAGCCTCTGCATCTTTTTTG-3’; shLKB1 5’-CCGGCGAAGAGAAGCAGAAAATGCTCGAGCATTTTCTGCTTCTCTTCGTTTTTG-3’; shAMPKα1 5’-CCGGGAGGAGAGCTATTTGATTACTCGAGTAATCAAATAGCTCTCCTCTTTTTG-3’;

### Viral Production and Transduction

Lentiviral particles were produced in 293FT cells according to the manufacturer’s instructions. HUVECs were grown to approximately 50% confluency. On the day of infection, cells were washed with 2x with PBS and incubated with viral supernatant containing 7µg/mL of polybrene for 15-18 h. After incubation, the cultures were washed, and full growth media was added. The infected cells were employed 48-72 hours after infection.

### Orbital Shear Stress

Orbital shear stress was applied to 90-95% confluent cells in growth media at the indicated amplitudes and durations, at 37°C and 5% CO_2_.

### Inhibitors and Functional Blocking Antibodies

Inhibitors were incubated with the cells 1 hour prior to application of orbital shear stress. CAMKKβ was inhibited using 10 µg/ml STO-609 (Selleck Chemicals). Myosin II was inhibited using 50μM Blebbistatin (Sigma). VE-Cadherin was inhibited using 4µg/ml BV9 (Invitrogen). GLUT-1 was inhibited using 30nM WZB117 (Tocris Biosciences). PFKFBP was inhibited using 25µM 3PO (Millipore).

### Immunoprecipitation and Immunoblotting

For immunoprecipitation of VE-Cadherin, cells were lysed in 1x RIPA (50 mM Tris-HCl pH 7.6, 150 mM NaCl, 1% NP-40, 0.5% Sodium Deoxycholate, 0.1% SDS, 50mM NaF, 2mM EDTA, 20μg/ml aprotinin, 2mM Sodium Vanadate, and 1mM PMSF). The lysates were clarified at 12,000 x g for 15 min at 4°C. The supernatant was collected, and 2 µg of VE-Cadherin (Cell Signaling) antibody was added and incubated for 2 hours at 4°C with rotation. The complexes were recovered using protein A beads, washed 3x in lysis buffer, and resuspended in 2x sample buffer (200 mM Tris, pH 6.8, 20% glycerol, 5% β-mercaptoethanol, 4% SDS, 0.3% Bromophenol Blue. Samples were boiled for 10 minutes and resolved using SDS-PAGE and transferred to PVDF (Immobilon). The membranes were blocked in 5% BSA for pAMPK, AMPK, pENOS, eNOS, VE-Cadherin or 5% Non-fat milk (Hyvee) for β-catenin, β-actin, and GAPDH. The membranes were incubated with the primary antibodies overnight at 4°C. Primary antibodies used for immunoblotting include: monoclonal Thr-172 phospho-AMPK at 1:1000 (Cell Signaling; 40H9), monoclonal AMPK-alpha at 1:1000 (Cell Signaling; 2532), monoclonal Ser-1177 phospho-eNOS at 1:500 (Cell Signaling; 9571S), monoclonal eNOS at 1:500 (Cell Signaling; D9A5L), monoclonal VE-Cadherin at 1:1000 (Cell Signaling; D87F2), monoclonal β-Actin at 1:1000 (Cell Signaling; 8H10D10), monoclonal β-Catenin at 1:1000 (BD Transduction; 610153), monoclonal LKB1 at 1:1000 (Cell Signaling; 27D10) or (Abcam; EPR19379) for coimmunoprecipitation studies. The membranes were washed 3x in TBST and blocked in either 5% BSA or 5% milk and incubated with anti-mouse HRP (Jackson Labs) or anti-rabbit HRP (Jackson Labs) at a 1:1000 dilution for 1 hour at room temperature. The membranes were washed 3x in TBST and visualized by using chemiluminescence detection reagent (Pierce), and signal was detected on an Odyssey Fc Imager (Li-Cor Biosciences). For analysis, the integrated density of each band was measured using the Image Studio Lite Ver 5.2 software. Ratios of the phosphorylated compared to the total proteins were quantified by stripping and re-probing membranes. Quantification of each assay represents a minimum of three experiments. Quantification was done using Studio Lite Ver 5.2 software and statistical analysis was conducted using GraphPad Prism 10.1.0 using either a two-tailed, unpaired student’s t-test when comparing two conditions, a one-way ANOVA with Dunnett’s comparison analysis when comparing more than two conditions, and a two-way ANOVA with Tukey’s analysis when comparing assays with multiple conditions and two variables.

### Immunofluorescence

Coverslips were coated with human fibronectin at 10 μg/ml, and cells were plated and allowed expand to confluency. Cells were fixed with 4% paraformaldehyde, permeabilized with 0.1% triton and washed with PBS. Cells were blocked with 10% BSA (Sigma) in PBS. The following reagents and primary antibodies were diluted in 10% BSA (Research Products International) and allowed to incubate O/N: Phalloidin conjugated with alexa-594 was used to visualize actin 1:1000, DAPI was used to visualize the nucleus 1:1000, monoclonal VE-Cadherin antibody at 1:500 (Invitrogen; 14-1449-82), monoclonal ENOS at 1:250 (Cell Signaling; D9A5L), and monoclonal Pecam-1 at 1:500 (Cell Signaling; 89C2). Cells were washed with PBS and the following secondary antibodies were diluted 1:1000 in 10% BSA anti-rabbit alexa-488 and/or anti-mouse alexa-647 and allowed to incubate for 2 hours at room temperature. Cells were washed with PBS and mounted on coverslips using MOWIAL (Sigma Corporation).

### Immunofluorescence quantification

Fluorescence images were captured at room temperature with a confocal microscope (model LSM 710; Carl Zeiss Micro Imaging, Inc.). A 40× or 60x oil objective (Carl Zeiss Micro Imaging, Inc.) with a numerical aperture of 1.3 was used. Images were obtained using the Zen2009 software (Carl Zeiss Micro Imaging, Inc.). Laser settings were kept identical for all conditions for each individual experiment. Quantifications of images were made using ImageJ. To determine enrichment of cellular junctions, the junctions of random cells were measured from at a minimum of three fields of view. Individual junctions were measured by creating ROI fit to the junction using the immunofluorescence of either VE-Cadherin for reference. Graphs report the intensity of the ROI. Pearson correlation coefficient was calculated by comparing two arrays of corrected fluorescence intensities for indicated number of junctions using the following equation: 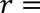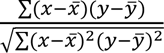

### Cellular Alignment

Cells were grown to 70-80% confluency on surfaces were coated with 10μg/ml of purified human fibronectin (SIGMA-Aldrich). Orbital shear stress was applied to the cells for 18-24 h at 37°C and 5% CO_2_. The cells were then washed 2x with PBS and fixed in 4% paraformaldehyde. Cells were washed and incubated in PBS containing 10% BSA containing 1:1000 Alexa-594 conjugated phalloidin and 1:1000 DAPI for 2-4 h at room temperature. Images were captured at room temperature with an inverted microscope (Axiovert 200M; Carl Zeiss), equipped with an ORCA-ERA 1394 HD camera (Hamamatsu Photonics, Hamamatsu City, Japan). A 10x Plan Neofluor objective (NA 0.55; Carl Zeiss) was used to capture fluorescent images.

### Assessing Endothelial Alignment

ImageJ v1.54 was used to calculate cellular alignment. Briefly, masks were created from the nuclear outline determined from the DAPI Image, by adaptive thresholding. The Analyze Particles feature was then used to quantify the parameters of the binary image. Here, a 0.0005-0.01 particle size threshold was used to remove nuclei that were overlapping or smaller artifacts. The Feret’s diameter was assessed and the Feret angle was used to assess cellular orientation. A cell was determined to be aligned to the direction of flow if the Feret angle was ±45° centered on the known direction of flow.

### Glucose Uptake Assay

Glucose uptake was assessed by measuring the fluorescence of 2-NBDG (Cayman Chemicals 11046). Cells were grown to confluency. Cells were pretreated with inhibitors or function blocking antibodies for 1 hour in glucose free media with 0.1mg/ml of 2-NBDG. Cells were then left under no shear or shear conditions for an additional hour. After incubation time the cells were washed 3x with PBS and lysed in EB lysis buffer (1 mM Tris-HCl, pH 7.6, 50 mM NaCl, 1% Triton X-100, 5 mM EDTA, 50 mM NaF, 20 μg/mL aprotinin, 2 mM Na3VO4, and 1 mM PMSF). Lysate was clarified by spinning down for 10 minutes at 12,000 rcf. Supernatant was transferred to 96 well plate and a fluorescence reading of 485/535 was taken (Biotek Synergy Neo model NEOALHPA B, Gen 5 software).

### Statistics and reproducibility

Statistical differences between groups of data were analyzed using a series of two-tailed unpaired Student t-tests for assays that contained only two conditions, a one-way ANOVA with Tukey’s analysis when comparing more than two conditions, and a two-way ANOVA with Tukey’s analysis when comparing assays with more multiple conditions and variables. All statistical analysis and data graphing was done on Prism (Version 10.1.0). All quantified experiments represent at least three biologically independent samples with the exception of immunofluorescence data. All immunofluorescence experiments were repeated independently three times with similar results.

