## Supplementary material for "Mechanical activation of VE-cadherin stimulates AMPK to increase endothelial cell metabolism and vasodilation": Supp. Figure 1

HUVECs

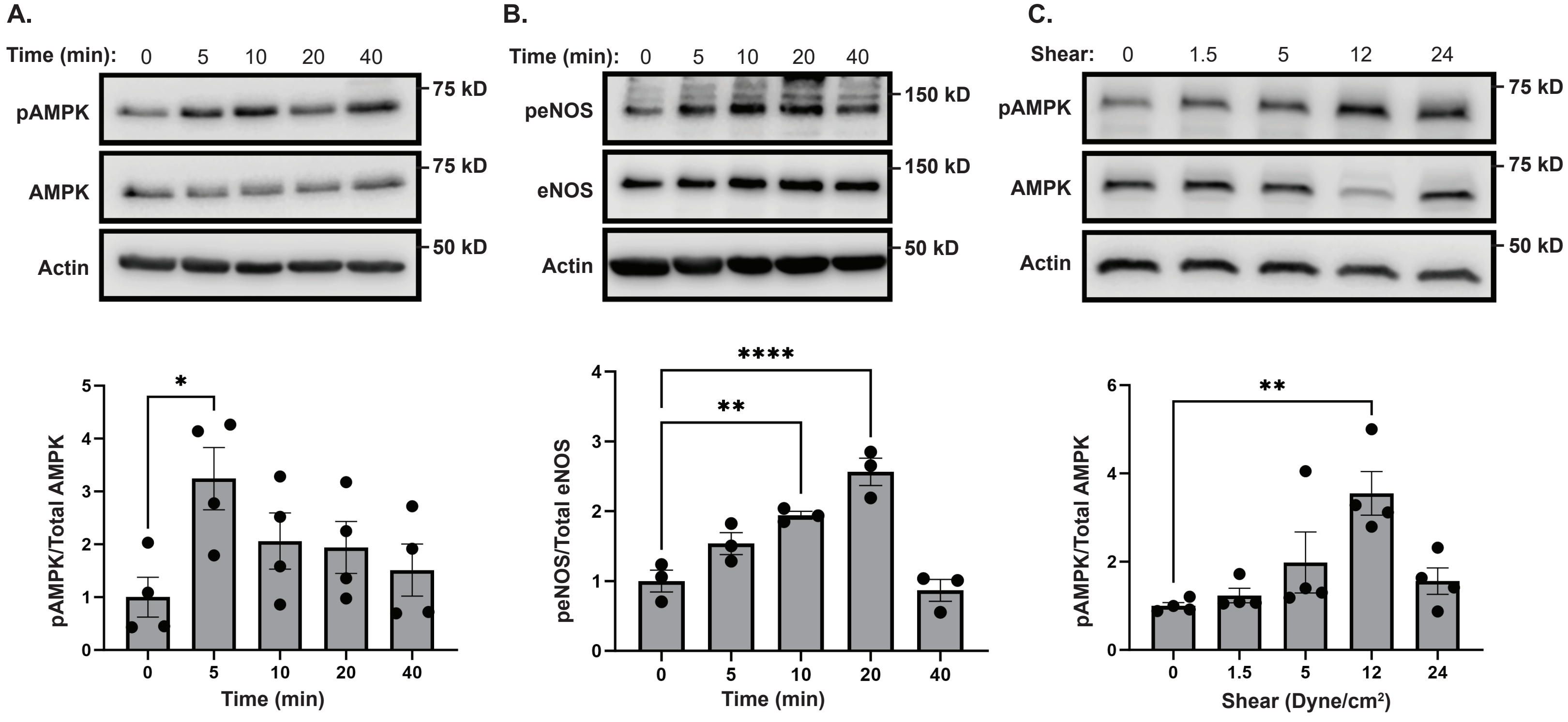


BAEC Data

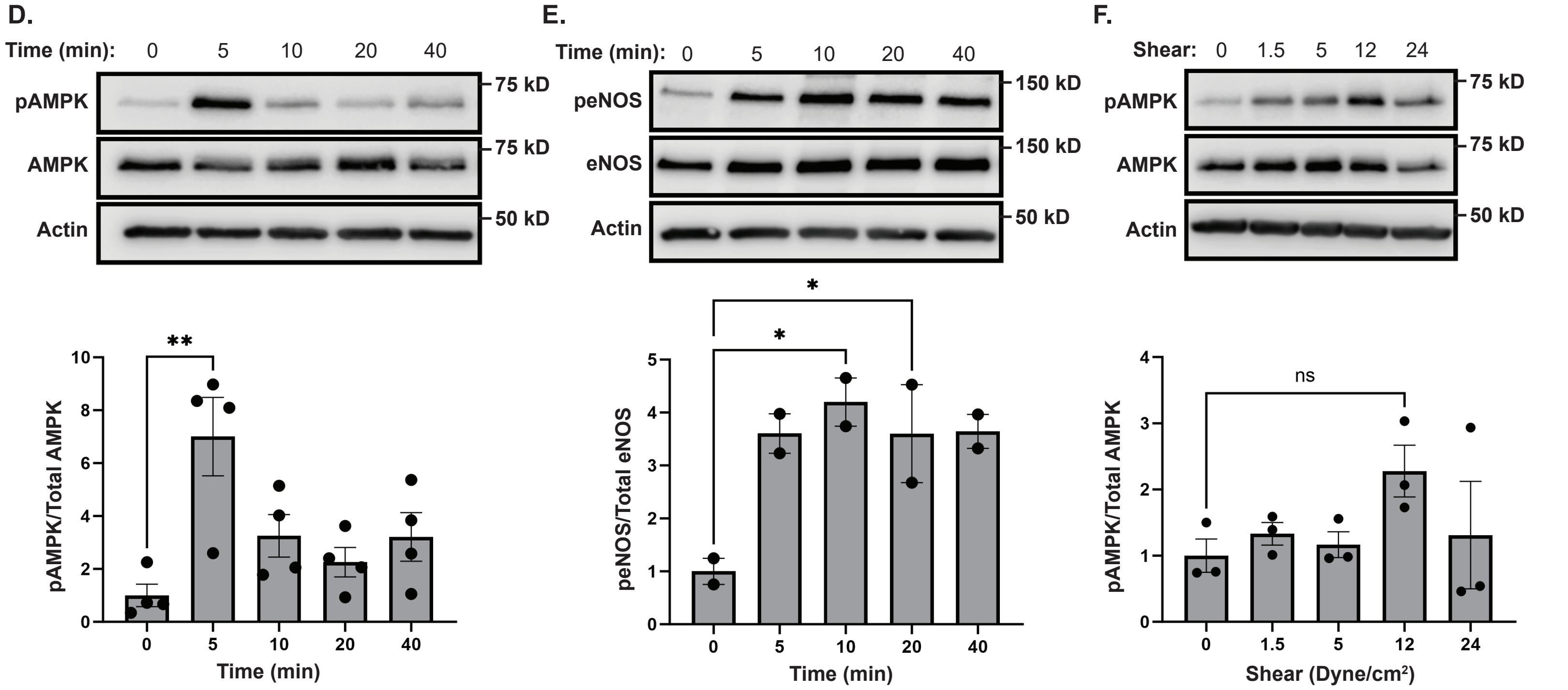


**Supplemental Figure 1:** (A-B) The time dependent activation of AMPK and eNOS in HUVECs. Cells were left exposed to shear stress for the indicated times. AMPK and eNOS activation were examined by immunoblotting total cell lysates with an antibody that recognizes AMPK phosphorylation in its activation loop (pAMPK) or phosphorylated Ser1177 on eNOS. Actin was blotted as a loading control. The blots were stripped and probed with antibodies that report on total AMPK levels (AMPK) or total eNOS levels (eNOS). The graphs below are the quantification of the ratios of phosphorylated AMPK to total AMPK and phosphorylated eNOS to total eNOS respectively; the data are mean ±s.e.m., n =4 or n=3 biologically independent samples. $*p<0.05$, $**p<0.01$, $****p<0.0001$ (one-way ANOVA with a Dunnett's comparison test). (C) The magnitude dependent activation of AMPK in HUVECs. Cells were left under no shear conditions or exposed to increasing magnitudes of shear for a constant time of 5 minutes. AMPK activation was examined by immunoblotting total cell lysates with an antibody that recognizes AMPK phosphorylation and actin was blotted as a loading control. The blots were stripped and probed for total AMPK. The graph below is the quantification of the ratios of phosphorylated AMPK to total AMPK; the data are mean ±s.e.m., n =4 biologically independent samples. $**p<0.01$, (one-way ANOVA with a Dunnett's comparison test). (D-E) BAECs were left exposed to shear stress for the indicated times. AMPK and eNOS activation were examined by immunoblotting total cell lysates with an antibody that recognizes AMPK phosphorylation in its activation loop (pAMPK) or phosphorylated Ser1177 on eNOS. Actin was blotted as a loading control. The blots were stripped and probed with antibodies that report on total AMPK levels (AMPK) or total eNOS levels (eNOS). The graphs below are the quantification of the ratios of phosphorylated AMPK to total AMPK and phosphorylated eNOS to total eNOS respectively; the data are mean ±s.e.m., n =4 or n=2 biologically independent samples. $*p<0.05$, $**p<0.01$, (one-way ANOVA with a Dunnett's comparison test), ns = not significant. (F) The magnitude dependent activation of AMPK in HUVECs. Cells were left under no shear conditions or exposed to increasing magnitudes of shear for a constant time of 5 minutes. AMPK activation was examined by immunoblotting total cell lysates with an antibody that recognizes AMPK phosphorylation and actin was blotted as a loading control. The blots were stripped and probed for total AMPK. The graph below is the quantification of the ratios of phosphorylated AMPK to total AMPK; the data are mean ±s.e.m., n =3 biologically independent samples. $**p<0.01$, (one-way ANOVA with a Dunnett's comparison test).
