## Supplementary material for "Mechanical activation of VE-cadherin stimulates AMPK to increase endothelial cell metabolism and vasodilation": Supp. Figure 2

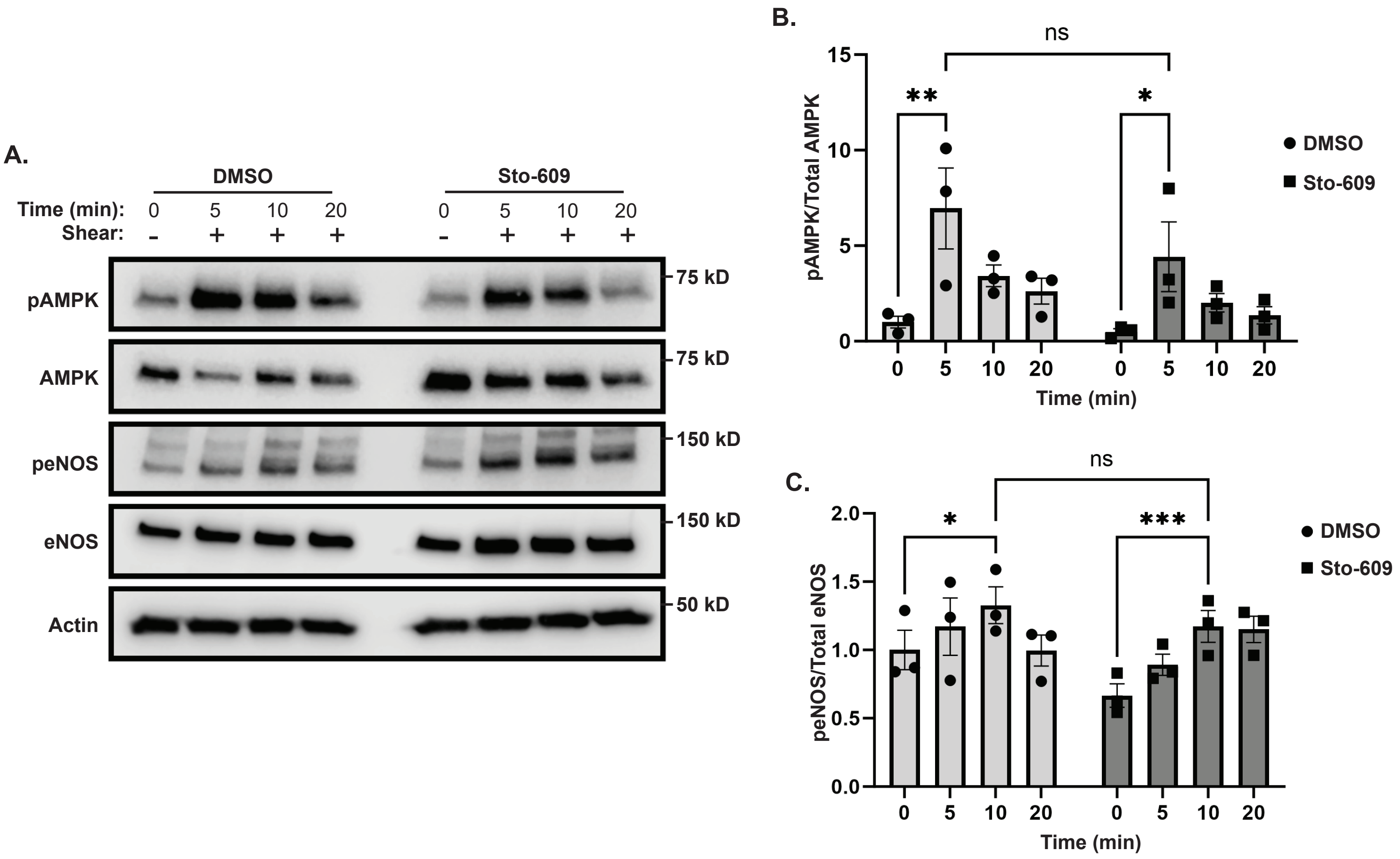


**Supplemental Figure 2:** (A-C). Shear stimulated AMPK and eNOS activation in the presence of the CaMKKβ inhibitor Sto-609. HUVECs were left resting (-) or exposed to shear stress (+) for the indicated times in the presence or absence of Sto-609 or DMSO as a control. AMPK and eNOS activation was examined by immunoblotting total cell lysates with an antibody that recognizes AMPK phosphorylation in its activation loop (pAMPK) or phosphorylated Ser1177 on eNO. Actin was blotted as a loading control. The blots were stripped and probed with antibodies that report on total AMPK (AMPK) or eNOS levels. The graph in B depicts the ratios of phosphorylated AMPK to total AMPK while the graph in C illustrates the ratios of phosphorylated eNOS to total eNOS; the data are mean ±s.e.m., n =3 biologically independent samples. *$p<0.05$, **$p<0.01$, ***$p<0.001$, (two-way ANOVA, with a Tukey's comparison test).
